# Seven SNPs in the Coding Sequence of Leptin Receptor Gene in Long-term Selected Japanese Quail Lines

**DOI:** 10.1101/044826

**Authors:** Kemal Karabağ, Sezai Alkan, Taki Karlı, Cengiz İkten, İnci Şahin, Murat Soner Balcıoğlu

## Abstract

The objective of this study was to identify SNPs in the coding sequence of the leptin receptor gene and to test for their possible association with 20 economically advantageous traits in 15 generations of 2 selected (HBW and LBW) and a control of japanase quail. A 350-bp part of the leptin receptor coding region was amplified and sequenced and understood that the fragment contained 7 SNPs (GenBank: KP674322.1-KP674328.1) that were detected in 5 loci (T3216C, T3265C, T3265G, C3265G, T3303C, A3311G, and T3347C) in a total of 30 individuals. The T3216C and T3303C SNPs located at the end of the codon were synonymous and did not affect the presence of proline. However, phenylalanine, leucine and valine were produced when the T3265C, T3265G and C3265G SNPs, respectively, were present. Glutamine or arginine was produced when the A3311G SNP was A or G, respectively, and serine was produced when the T3347C SNP was C. Although codons and amino acid sequences changed due to the second SNP, the secondary protein structure was not changed. However, the fourth and fifth SNPs changed both the amino acid sequences and secondary protein structure.

Pairing the SNP loci with phenotypic traits created haplogroups. When all individuals were evaluated together, some of the differences between the haplogroups were statistically significant (p<0.05; p<0.01). These results showed that both the sequence and structure of the leptin receptor gene could be altered by long-term selection. However, to achieve a more precise understanding of the role of leptin, entire coding sequences of leptin and the leptin receptor should be studied.

## INTRODUCTION

The control of feed intake, energy balance and fat deposition has high economic importance in farm animals. The accumulation of excess fat affects meat quality, fertility, productivity, and whole body metabolism (Macajova et al. 2004). Leptin activity at specific receptors in the hypothalamus suppresses feed intake, which increases the use of energy, and leptin is a polypeptide hormone that controls the body’s energy balance (Taouis et al. 1998; Ashwell et al. 1999). Leptin stimulates liposis, insulin sensitivity, glucose utilization in muscle, and fatty acid oxidation in liver, muscle and adipose tissue (Schwartz et al. 1996; Ahima and Osei 2004).

The leptin gene was first cloned and sequenced from chicken in 1998 (Taouis et al. 1998; Ashwell et al. 1999). DNA sequence analysis of leptin from several mammalian and avian species, including livestock animals, showed that leptin is highly conserved among vertebrate species. Among vertebrates, the leptin amino acid sequence shows approximately 80% homology (Doyon et al. 2001). Wang et al. (2014) sequenced the entire coding region of the leptin receptor in *C. coturnix japonica*. The presence of leptin in laying and broiler poultry genotypes decreases feed intake (Denbow et al. 2000; Kuo et al. 2005). Quail chicks that were given recombinant mouse leptin on the fifth embryonic day were removed from incubation earlier (5-24 hours) and reached a higher body weight than the control group (Lamosova et al. 2003). In male quails that were given leptin during the embryonic period, the testes were two-fold larger, and testosterone levels were higher compared to those of controls on the 35^th^ day. Moreover, female individuals who were treated with leptin began laying eggs earlier, and the number of eggs from each individual was higher than that of the control group (Macajova et al. 2002).

Mutations in genes responsible for the biochemical activity of an organism, such as those in leptin, can result in quantitative differences. With advances in DNA sequencing methods and technologies, molecular studies in recent years, including those in livestock species, have focused on single nucleotide polymorphisms (SNPs). In parallel, determining the DNA sequences and SNPs of leptin and its receptors in relation to livestock yields has accelerated research efforts. Researchers are actively investigating the relationship between leptin and traits of high economic importance, such as feed intake (Lagonigro et al. 2003; Huang et al. 2011), fertility (Liefers et al. 2002), milk production (Banos et al. 2008; Chebel and Santos 2011) and meat production (Lusk 2007; Boucher et al. 2011).

In this study, possibility of SNPs that might be occurred in long-term selection in the coding sequences of leptin receptor gene and their relationship with phenotypic traits were investigated in 15 generations of selected Japanese quail (*Coturnix coturnix japonica*).

## MATERIALS AND METHODS

In this study, three different populations of Japanese quail were used as the research model. One population was used as the control (C), which was not selected previously. The other two populations were extremely different from each other because they were selected for 15 generations for high body weight (HBW) and low body weight (LBW) in two different previous projects (Project numbers 21.01.0121.30 and 2003.03.121.004) that were performed at the Scientific Research Projects Coordination Unit of Akdeniz University.

### Collection of Eggs and Chick Rearing

Fertilized eggs were collected from the HBW, LBW and C populations for a week and stored at 15-20 °C and 75-80% humidity. These eggs were incubated at 36.5 °C and 65% humidity for the first 14 days and at 36.0 °C and 55% humidity for the last 4 days. Chicks were weighed individually using 0.01 g precision scales, and an aluminum ID number was attached to the left wings of chicks after incubation. These chicks (40 birds) were fed 24% crude protein and 2900 kcal/kg ME (metabolic energy) during the first four weeks in a breeding cage. Sex determination was performed by observing the cloaca and breast feather color at the end of the fourth week. Males and females (n=50 for both) were selected randomly from each population and transferred to individual breeding cages for 10 weeks. All birds were fed 21% crude protein and 2800 kcal/kg ME for ten weeks. Lighting was applied continuously for the first four weeks and then 16 hours a day.

### Body Weight and Feed Intake

Body weights (B1W) were measured for each animal using 0.01 g precision scales from hatching (H1W) to the 15^th^ week on the same day each week. The differences arising from feed intake (FI) were fixed because quails were not fed on the day of weighing. Thus, 10 weeks of individual feed intake (FI) were recorded as grams per day.

### Sexual Maturity

The first ovulation day and the first day of release foam were considered as sexual maturity for females and males, respectively. Females and males were weighed on the same day. In this manner, sexual maturity age (SMA) and sexual maturity weight (SMW) were determined.

### Egg Yields of Females

Egg yields (number and weight) of female quails that were transferred from individual breeding cages were recorded daily for 10 weeks. The total egg weight (TEW) and total egg number (TEN) were obtained.

### Carcass Traits

A total of 30 quails selected randomly as 5 females and 5 males from the HBW, LBW and C populations were introduced to the cutting process after five weeks of growth and yield during a ten-week period. The body weights of the selected animals were measured just before cutting. Low-voltage electrical current (100 mA, 50 Hz) was used to stun animals as recommended in the relevant scientific literature (Göksoy et al. 1999), and then the jugular vein was cut. Separation of the carcasses of the slaughtered quails was performed according to the recommendations in the literature (Yalçın et al. 1995). The body weight (B1W), breast weight (B2W), back weight (B3W), carcass weight (CW), gizzard-null weight (GW), head weight (H2W), heart weight (H3W), liver weight (LW), left leg weight (LLW), left wing weight (LWW), right leg weight (RLW) and right wing weight (RWW) were measured using 0.01 g precision scales. The measurements of the carcass components were obtained immediately after cutting. Tibia bone length (TBL) and tibia bone width (TBW) were measured using a digital compass after cleaning and separating the surrounding body tissue. Bone lengths were measured from the proximal to distal ends of the bones.

### Tissue Samples and Total RNA Extraction

Liver tissue samples from each individual were isolated using sterile forceps and scissors immediately after cutting and placed into numbered tubes (Corning, New York, U.S.A.) containing RNAsave. These tissues were stored at -80 °C until use. Cellular degradation of the liver tissues was performed using a lyser with tungsten beads. Then, total RNA extraction was performed using a commercial kit (Axygen). The total RNA obtained from 30 quails was stored at –80 °C until use.

### cDNA Synthesis and PCR Amplification

A commercial kit (Thermo Scientific #K1621) was used to synthesize cDNA from total RNA using the following protocol: 60 min at 42 °C, 5 min at 70 °C. Primers (forward, gcttgctcaggtagctcctg and reverse, tgcggcacgtatggcacgat) based on the recommendations of Dridi et al. (2005) were used to PCR amplify a 350 bp leptin receptor coding region from cDNA. PCR products (15 µl) were evaluated for a 350 bp length using 2% agarose gels (electrophoresed at 80 V/2 h) and stained with ethidium bromide. Separated fragments in the electrophoresis gel were cut with a sterile scalpel under UV light and transferred to individual 1.5 ml pre-numbered tubes. The PCR reactions were performed in 20 ml reaction volumes with 2 µl of genomic DNA (20 ng) as a template, 2 µl of buffer (NH_2_SO_4_), 0.4 µl of a dNTP mix (2.5 mmol/L), 0.5 ml of each primer (20 nmol/ml), 1.25 µl of MgCl_2_ (25 mM) and 0.15 µl of EX Taq polymerase (Takara Bio Inc. Shiga, Japan). Amplifications were performed using a thermal cycler (Thermo Arktik) with the following conditions: 3 min for an initial denaturation at 94 °C, 35 cycles at 94 °C for 30 s for denaturation, 30 s for annealing at 60 or 62 °C, 45 s for extension at 72 °C, and a final extension for 5 min at 72 °C. *β-actin* gene primers (forward, *caaggagaagctgtgctacgtgc* and reverse, *ttaatcctgagtcaagcgcc*) were used to determine that the PCR protocol worked.

### Sequence Analysis, SNP Determination and Haplogroup Generation

DNA samples were concentrated in the PCR and sequenced directly in a sequencing instrument (Beckman 8800) after being purified from a gel and denatured at 94 °C. A leptin receptor gene fragment of 350 bp was sequenced for 30 individuals from the HBW, LBW and C genotypes. These sequences were compared with each other for the presence of SNPs. Seven SNPs were found in five loci. Haplogroups were created by pairing the SNP locus and phenotypic traits and formed by grouping SNPs both between and within genotypes.

### Statistical Analysis

To determine the location of the DNA fragment (350 bp) in the leptin receptor gene, the amino acid sequence of the DNA fragment and the secondary protein structures from these amino acid sequences were determined using the online software at http://blast.ncbi.nlm.nih.gov/Blast.cgi, http://translate-protein.com/ and http://www.biogem.org/. The predicted amino acid changes were located, and the significant differences were investigated.

All phenotypic traits (B1W, B2W, B3W, H1W, H2W, H3W, RWW, LWW, RLW, LLW, LW, CW, GW, SMW, TEW, SMA, TEN, FI, TBW and TBL) were evaluated using a Kolmogorov-Smirnov test for the normal distribution of all the data from three groups together and each group separately. A Box-Cox transformation was applied to the trait data that was not normally distributed, and the converted data were tested again for normality. B1W, LW, SMW, TBL and TBWA showed a normal distribution, whereas other features did not. When the normality test was applied to genotypes separately, several traits were not normally distributed, including FI, GW, H1W, LW, LLW, LWW, RLW, SMA, TBW in LBW; CW, GW, H3W, SMA, TBL in HBW; GW, H1W, H3W, SMA in control.

A t-test and Mann Whitney U test were conducted for binary SNPs (3216, 3303, 3311, and 3347) and haplogroups, respectively, and for traits that showed either a normal distribution or a non-normal distribution, respectively, in individuals who clustered in the same population but with different SNP haplogroups. ANOVA and Kruskal Wallis tests were applied for normally distributed data and non-normally distributed data, respectively, which had three alternative haplogroups (3265). In addition, the **χ**2 test statistic was used on each of the SNPs to determine whether the ratio was due to genotype possession.

### One-way ANOVA Procedure

*Model: y_ij_ = μ + g_i_ + e_ij_*

*Where;*

*y:* traits; *g_i_*: i. SNP effect; and *e_ij_*: error.

## RESULTS AND DISCUSSION

A 350-bp fragment of the leptin receptor gene was sequenced in 30 individuals from three genotypes (Table 1). These fragments were BLAST searched against GenBank to confirm their identity as fragments of the leptin receptor gene *(Coturnix coturnix japonica;* KJ639903.1). A 116 amino acid sequence is predicted for the 350-bp fragment (Table 1). The fragment was compared among 30 individuals, and seven SNPs were identified at five loci (Table 2). The sequences and SNPs were published in the GenBank website with the accession codes KP674322.1-KP674328.1 (http://www.ncbi.nlm.nih.gov/nuccore/).

**Table 1.**
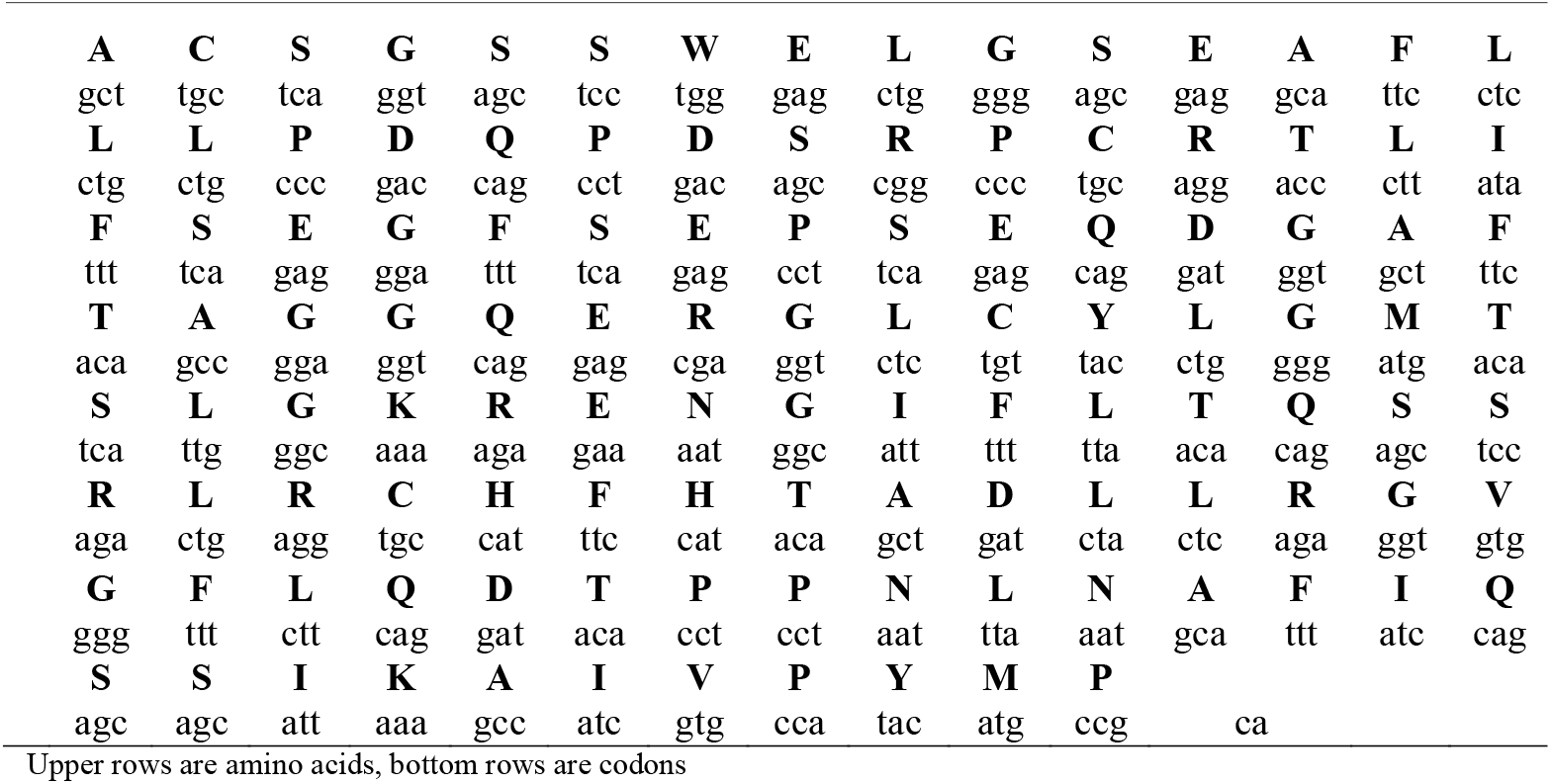
A 350 bp part of the leptin receptor gene and its amino acid sequence are shown.

**Table 2.** The SNPs identified in 30 individuals of the C, HBW and LBW genotypes.

| Genotypes | SNP locus | individuals |  |  |  |  |  |  |  |  |  |
| --- | --- | --- | --- | --- | --- | --- | --- | --- | --- | --- | --- |
|  |  | 1 | 2 | 3 | 4 | 5 | 6 | 7 | 8 | 9 | 10 |
| C | 3216 | t | c | t | t | c | c | c | c | c | t |
|  | 3265 | t | t | t | t | t | t | t | t | t | c |
|  | 3303 | c | c | t | t | c | c | c | c | c | t |
|  | 3311 | a | a | g | g | a | a | a | a | a | g |
|  | 3347 | t | t | t | t | t | t | t | t | t | t |
| HBW | 3216 | c | t | c | t | t | c | t | c | t | t |
|  | 3265 | g | g | g | g | g | g | g | g | g | g |
|  | 3303 | c | t | c | t | t | c | t | c | t | t |
|  | 3311 | g | g | a | g | g | a | g | a | g | g |
|  | 3347 | c | t | c | t | t | t | t | t | t | t |
| LBW | 3216 | c | c | c | c | c | c | c | c | c | c |
|  | 3265 | c | c | t | c | c | c | c | t | c | t |
|  | 3303 | t | c | c | c | c | t | t | c | c | c |
|  | 3311 | g | g | a | g | g | g | g | a | a | a |
|  | 3347 | t | t | t | t | t | t | t | t | t | t |

C: control, HBW: High Body Weight, LBW: Low Body Weight

Seven DNA sequences with seven SNP loci were uploaded to GenBank (http://www.ncbi.nlm.nih.gov/nuccore/KP674322) under the accession numbers KP674322-328. Wang et al. 2014 uploaded the sequence of the entire leptin receptor mRNA (3579 bp) of *Coturnix coturnix japonica* to GenBank under the accession number KJ639903.1 (http://www.ncbi.nlm.nih.gov/nuccore/KJ639903.1). The 350 bp part of the sequence that we identified exactly matched bases from 3163 to 3513 of the 3579 bp leptin receptor sequence.

The codons and amino acid sequence of the leptin receptor were determined using the online program http://translate-protein.com (Table 1.). Other possible codons and amino acid sequences were determined for other SNPs of this DNA sequence. The other possible codons and amino acid sequences that were derived from these SNP loci were different from each other. Changes in the secondary protein structures due to the SNPs were determined using an online program (http://www.biogem.org).

There was no amino acid difference when the first and third SNP were C and T because these SNPs only changed the last nucleotide of the codon. However, phenylalanine (F), leucine (L) and valine (V) were formed when the second SNP was T, C and G, respectively. Glutamine (Q) and arginine (R) were formed when the fourth SNP was A and G, respectively. Leucine (L) and serine (S) were formed when fifth SNP was T and C, respectively.

Structural changes in the amino acid sequence and protein were examined to evaluate the functional effects of the SNPs. From the 350-bp fragment, 116 amino acids were formed. The secondary protein structure of leptin receptor according to the basic amino acid sequence showed 63.8% α-helices, 59.5% β-sheets, 17.2% turns and 14.7% coils. The amino acid sequence was not altered because the first and third SNPs were found in the third nucleotide of the codons in question and thus resulted in no changes to the protein structure. The seven possible amino acid sequences were published in GenBank (http://www.ncbi.nlm.nih.gov/protein/) under the accession codes AKM16812.1-AKM16818.1.

Although codons and amino acid sequences changed due to the second SNP, the secondary protein structure was not changed. However, the fourth and fifth SNPs changed both the amino acid sequences and secondary protein structure. When the fourth SNP occurred, there were 3.8% α-helices, 53.4% β-sheets, 17.2% turns, and 19.0% coils, and for the fifth SNP, there were 62.9% α-helices, 57.8% β-sheets, 17.2% turns and 17.2% coils.

When a t-test and Mann-Whitney U test were performed for traits that have a normal distribution (B1W, LW, SMW, TBL and TBW) and not-normal distribution (B2W, B3W, CW, FI, GW, H1W, H2W, H3W, LLW, LWW, RLW, RWW and SMA), respectively, for SNP 1, statistically significant differences (p>0.05, p>0.01) were found between SNP haplogroups, with the exception of LW, FI and G (Tables 3, 5).

**Table 3.** A t-test of the SNP haplogroups for the normally distributed phenotypic values for all individuals 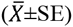.

| SNP loci | alleles | N | BW1 | LW | SMW | TBL | TBW |
| --- | --- | --- | --- | --- | --- | --- | --- |
| 3216 | c | 20 | 207.27±17.78* | 3.82±0.45 | 188.70±16.20* | 47.39±1.26** | 2.21±0.18** |
| | t | 10 | 278.88±16.99* | 4.31±0.50 | 257.46±20.11* | 52.83±1.05* | 2.99±0.17* |
| 3303 | c | 18 | 218.00±18.53 | 3.99±0.48 | 197.97±17.00 | 48.11±1.35 | 2.28±0.19 |
| | t | 12 | 250.85±22.60 | 3.96±0.49 | 232.10±23.16 | 50.85±1.49 | 2.76±0.21 |
| 3311 | g | 16 | 235.50±22.43 | 3.61±0.45 | 214.14±21.55 | 49.15±1.52 | 2.50±0.23 |
| | a | 14 | 226.16±18.00 | 4.41±0.51 | 208.74±17.53 | 49.26±1.38 | 2.44±0.17 |
| 3347 | c | 2 | 348.12±18.48* | 5.06±0.46 | 297.36±0.35 | 55.82±2.12 | 3.34±0.06 |
| | t | 28 | 222.78±14.10* | 3.90±0.36 | 205.50±14.17 | 48.73±1.03 | 2.41±0.15 |

\* and \*\* with different letters differ significantly at $P<0.05$ and $P<0.01$ , respectively.
BW1: Body Weight, LW: Liver Weight, SMW: Sexual Maturity Weight, TBL: Tibia Bone Length, TBW: Tibia Bone Weight,

**Table 4.**
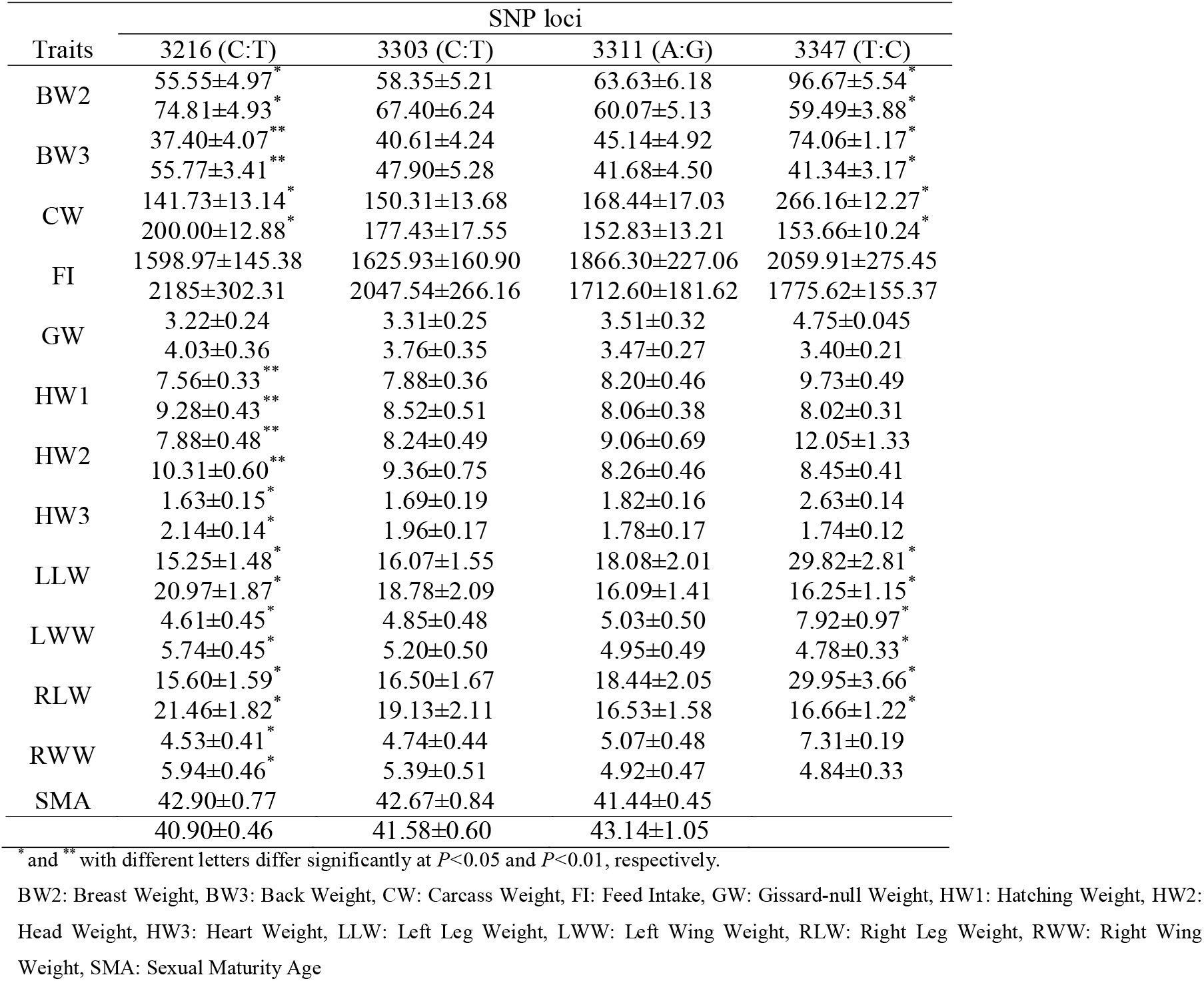
Mann-Whitney U test of SNP haplogroups for the phenotypic values from all individuals 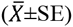.

**Table 5.** ANOVA test of SNP haplogroups that have three alleles at the 3,265 (SNP 2) locus for normal distribution phenotypic values from all individuals 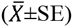.

| alleles | traits |  |  |  |  |
| --- | --- | --- | --- | --- | --- |
|  | BW1** | LW** | SMW** | TBL** | TBW** |
| c | 150.57±12.10 | 2.47±0.42 | 138.30±12.85 | 43.42±1.11 | 1.73±0.21 |
| t | 206.43±11.38 | 4.34±0.59 | 185.45±10.77 | 47.63±0.79 | 2.26±0.15 |
| g | 325.25±8.96 | 4.77±0.49 | 301.69±10.91 | 55.72±0.59 | 3.33±0.08 |

\*\* with different letters differ significantly at $P<0.01$ .
BW1: Body Weight, LW: Liver Weight, SMW: Sexual Maturity Weight, TBL: Tibia Bone Length, TBW: Tibia Bone Weight

Additionally, when an ANOVA and Kruskal-Wallis test were performed for traits that are normally distributed (B1W, LW, SMW, TBL VE TBW) and not-normal distribution (B2W, B3W, CW, FI, GW, H1W, H2W, H3W, LLW, LWW, RLW, RWW and SMA) in SNP 2, statistically significant differences (p>0.05) were found between SNP haplogroups, with the exception of SMA (Tables 5, 7). However, when a t-test and a Mann-Whitney U test were performed for traits that have a normal distribution (B1W, LW, SMW, TBL and TBW) and non-normal distribution (B2W, B3W, CW, FI, GW, H1W, H2W, H3W, LLW, LWW, RLW, RWW and SMA) for SNP 3 and SNP 4, there were no statistically significant differences. On the other hand, significant differences were found for the normally distributed B1W trait in SNP 5 and the non-normally distributed B2W, B3W, CW, LLW, LWW, and RLW traits (p>0.05). There was no difference between the haplogroups in terms of studied phenotypic characteristics in the control vs. LBW genotypes. A statistically significant difference was found between haplogroups H3W and CW in SNP 4 and SNP 5, respectively, and in the HBW genotype (p<0.05).

**Table 6.**
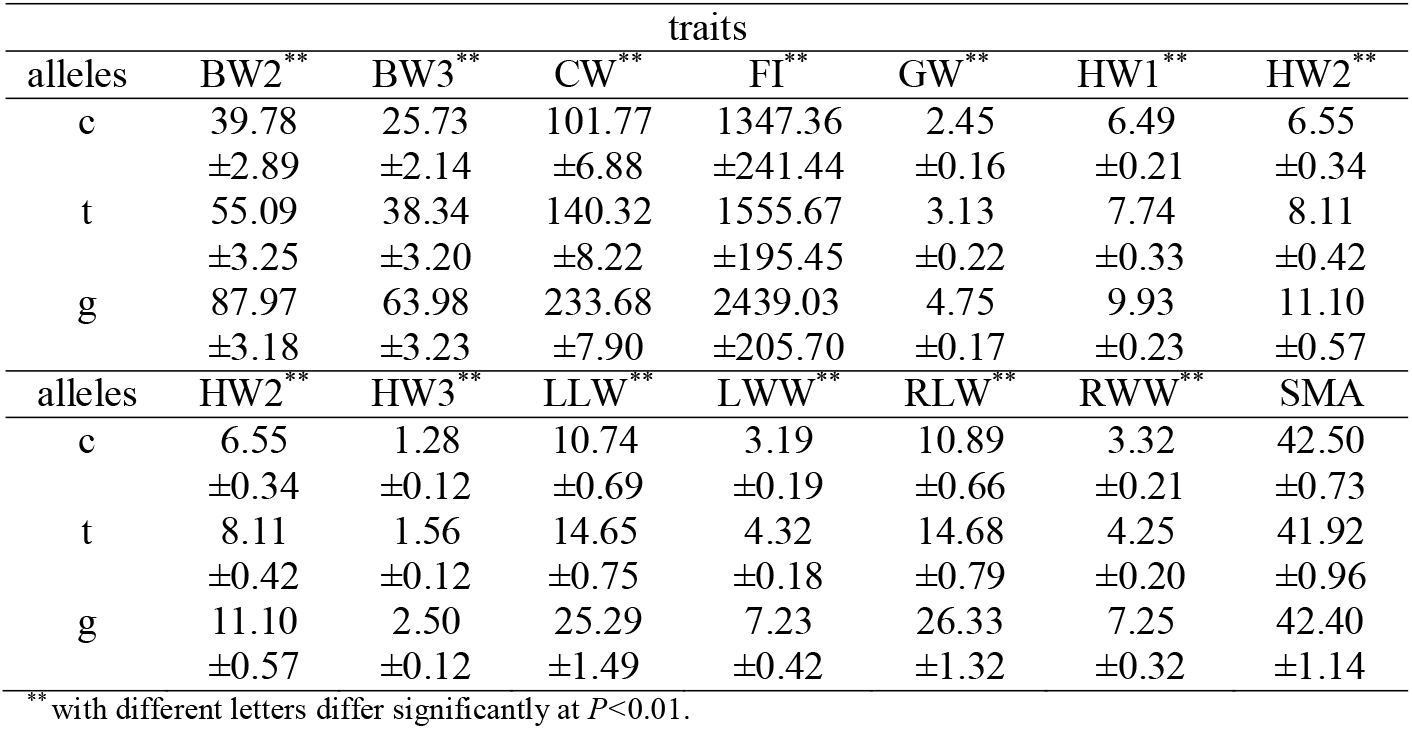
Kruskal-Wallis test of SNP haplogroups that have three alleles at the 3,265 (SNP 2) locus for non-normal distribution phenotypic values from all individuals 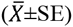.

However, no statistically significant differences in the SNP haplogroups from the phenotypic parameters of the non-normal distribution were found when the genotypes were tested separately.

Additionally, a χ^2^ test was performed to determine whether the SNP ratio depended on genotypes, according to the calculations of SNPs 1 and 2. The results showed that the SNPs were due to the genotype of the SNP ratio (p<0.05).

All statistical analyses suggested that SNPs 1 and 2 altered the function of the leptin receptor. However, conclusively demonstrating this effect requires the identification of all of the SNPs in the entire sequence of the leptin receptor in quails.

Although leptin function is controversial in chickens (Sharp et al. 2008), leptin is an excellent candidate gene for livestock production, as it is associated with features of economic importance. Indeed, in recent years, leptin gene polymorphism studies of several single nucleotide polymorphisms (SNPs) have been identified in cows and pigs (Jiang and Gibson 1999; Buchanan et al. 2002; Liefers et al. 2002; Buchanan et al. 2003; `Lagonigro et al.` 2003; Lusk 2007). Several SNPs have been identified that are associated with important economic traits, such as milk yield, feed intake, adiposity, growth, and carcass quality. DeVuyst et al. (2008) found that both crossbred CT and TT beef cows wean significantly heavier beef calves than CC cows. There were several SNPs found in the porcine *LEP* and *LEPR* genes, suggesting that the SNPs lead to increased growth and fat (Perez-Montarelo et al. 2012). Buchanan et al. (2003) revealed a SNP in the leptin gene of dairy cattle. This polymorphism, in which the first nucleotide is a thymine instead of cytosine in the 25^th^ codon, changed arginine to a cysteine. However, homozygous animals carrying the T allele show no difference compared to animals carrying the C allele in terms of milk fat and milk protein production and a daily milk yield of more than 1.5 kg. In addition, homozygous T allele-bearing animals developed higher fatty carcasses than those with the C allele (Buchanan et al. 2002). Leptin gene polymorphisms are used as DNA markers in marker-assisted selection (MAS) for commercial farming.

Feed consumption, growth, development, energy metabolism and immune system functioning have a high economic importance in livestock. In this regard, there is a need for additional studies on genes that affect animal feed intake, the regulation of energy metabolism, yield and health. Leptin plays an important role in all of these mechanisms of economic importance in livestock. Therefore, studies of leptin will significantly contribute to animal nutrition, breeding and health.

The effects of long-term selection in reared quail genotypes on the leptin receptor depend on the presence of specific SNPs, and these SNPs result in significant changes in the secondary structure of proteins. However, to achieve a more accurate understanding of the role of leptin and its receptor, the DNA sequence of all of the SNP changes that benefit individuals and alter protein structure should be identified.

## ACKNOWLEDGEMENT

This study was supported by the Akdeniz University, The Scientific Research Projects Coordination Unit (Antalya) with project Number 2012.01.0104.002.

All the experimental procedures were reviewed and approved by Akdeniz University Local Committee on Animal Research Ethics. Protocol number is 2012.02.02.

